# Evidence for a conserved queen-worker genetic toolkit across slave-making ants and their ant hosts

**DOI:** 10.1101/2021.10.20.465091

**Authors:** B. Feldmeyer, C. Gstöttl, J. Wallner, E. Jongepier, A. Séguret, D.A. Grasso, E. Bornberg-Bauer, S. Foitzik, J. Heinze

## Abstract

The ecological success of social Hymenoptera (ants, bees, wasps) depends on the division of labour between the queen and workers. Each caste is highly specialised in its respective function in morphology, behaviour and life-history traits, such as lifespan and fecundity. Despite strong defences against alien intruders, insect societies are vulnerable to social parasites, such as workerless inquilines or slave-making (dulotic) ants. Here, we investigate whether gene expression varies in parallel ways between lifestyles (slave-making versus host ants) across five independent origins of ant slavery in the *“Formicoxenus*-group” of the ant tribe Crematogastrini. As caste differences are often less pronounced in slave-making ants than non-parasitic ants, we also compare the transcriptomes of queens and workers in these species. We demonstrate a substantial overlap in expression differences between queens and workers across taxa, irrespective of lifestyle. Caste affects the transcriptomes much more profoundly than lifestyle, as indicated by 37 times more genes being linked to caste than to lifestyle and by multiple caste-associated gene modules with strong connectivity. However, several genes and one gene module are linked to the slave-making lifestyle across the independent origins, pointing to some evolutionary convergence. Finally, we do not find evidence for an interaction between caste and lifestyle, indicating that caste differences remain consistent even when species switch to a parasitic lifestyle. Our findings are a strong indication for the existence of a core set of genes whose expression is linked to the queen and worker caste in this ant taxon, supporting the “genetic toolkit” hypothesis.

## Introduction

The ecological success of social insects is based on the efficient division of labour between reproductives and non-reproductives, i.e., in the social Hymenoptera, the queens and workers (Hölldobler & Wilson, 2009; Wilson, 1971). Instead of producing their own offspring, workers help to raise the offspring of their mother or other related queens. The altruism of workers is explained by their relatedness to the recipients of their help, which is maintained by the closure of the society against unrelated freeloaders or parasites (Hamilton, 1964, 1987). Nevertheless, several species have evolved sophisticated ways to infiltrate and usurp social insect colonies (Rabeling, 2020), and among these are the charismatic “slave-making” or “dulotic” ants (Hölldobler & Wilson, 1990; Buschinger, 2009; D’Ettorre & Heinze, 2001; Mori et al., 2001; Visicchio et al., 2001). Freshly mated, young queens of slave-making ants invade the nests of closely related, non-parasitic ant species, where they kill or expel the resident queen(s) and often also the adult workers. Host workers emerging from the conquered host brood take care of the slave-maker queen and her offspring, maintain the nest, and forage for food. Slave-maker workers do not engage in normal worker chores (Hölldobler & Wilson 1990; Buschinger, 2009) but instead raid neighbouring host nests and pillage their brood, thus replenishing or increasing the host force.

Ten independent origins of slavery in ants are documented (Stoldt & Foitzik, 2021), and at least five convergent origins lie within the “*Formicoxenus*-group” of the myrmicine tribe Crematogastrini (Blaimer et al., 2018). This allows investigation of whether convergent changes in gene expression occurred among those evolutionary switches to slave-making. There are several morphological, physiological and behavioural similarities among slave-making species of independent origin. For example, slave-maker workers are often heavily armed with strong mandibles and associated muscles, which leads to enlarged heads (Hölldobler & Wilson, 1990). Given that workers of slave-making species no longer take care of the daily duties in the colony, they have become more queen-like in their task repertoire. They neither forage for food nor do they engage in brood care (Buschinger, 2009). Moreover, slave-maker workers tend to have increased reproductive potential.

While the ovaries of non-parasitic workers have fewer ovarioles than the queen’s ovaries and rarely contain mature eggs in the presence of a fertile queen, the ovaries of slave-maker workers often have the same number of ovarioles as the queen (Heinze, 1996b). They form reproductive hierarchies (Franks and Scovell, 1983; Bourke, 1988; Heinze, 1996a) and frequently lay male-destined eggs even in the queen’s presence (Foitzik & Herbers, 2001; Brunner et al., 2005; Suefuji & Heinze, 2014). Gene expression differs strongly between workers and queens of most species, reflecting their divergent function, behaviour, and physiology (Gstöttl et al., 2020; Korb et al., 2021; Morandin et al., 2019a, b; Feldmeyer et al., 2013). For example, queen transcriptomes are characterised by the expression of genes associated with fecundity (e.g. vitellogenins), immunity, DNA repair and response to oxidative stress functionalities linked to their long lifespan (Stoldt et al., 2021). However, given the described similarities between slave-maker queens and workers, we expected their transcriptomes to differ less than those of queens and workers of non-parasitic species.

We therefore investigated the influence of caste (queen vs worker) and lifestyle (slave-maker vs non-parasitic) on gene expression of adult individuals and the interaction between those two parameters, which would indicate that caste is affecting gene expression differently in non-parasitic versus slave-making species. Here we use a protocol which is aimed at maximising reproducibility and minimising confounding effects by using all five available and evolutionary closely related pairs of slave-maker and host in the myrmicine tribe Crematogastrini. For each species, we sequenced the transcriptomes of six pooled workers and three pooled queens, respectively, taking slave-making species vs. host species as replicates. We investigated gene regulatory network properties according to caste and lifestyle and constructed orthologue clusters to investigate putative parallel selection patterns in genes associated with the slave-maker versus host lifestyle, including two related non-host taxa and two samples of the distantly related ant *Cardiocondyla obscurior* as outgroup.

## Material & Methods

### Sampling and sequencing

Colonies of 15 myrmicine ant species of the “*Formicoxenus*-group” (genera *Harpagoxenus, Leptothorax*, and *Temnothorax*, including the previously synonymized genera of slave-making ants: *Chalepoxenus, Myrmoxenus*, and *Protomognathus* (Ward et al., 2015, but see Seifert et al., 2016) were collected between 2016-2018 from various locations across Germany, Italy, and the US (Supplement_coordinates). Colonies were either brought to the lab in Regensburg, Mainz, or Münster, and kept under standard conditions in incubators (12 h 25°/ 12 h 25°C day-night cycles) before six workers and three queens were pooled per species for RNA extraction. For each species, workers and queens originated from three colonies. We generated one queen and one worker transcriptome for seven slave-making ants and their host species (Supplement Table S1). RNA was extracted using the Nucleo-Spin Mini kit (Macherey-Nagel). Samples were shipped to StarSEQ (Mainz) for library preparation and 100bp paired end sequencing on an Illumina HiSeq. In total, we obtained 14mio reads on average per sample (Supplement Table S2).

### Gene expression analyses

Raw reads were quality checked using FastQC v.0.11.8 (Andrews, 2010), and adapters trimmed with Trimmomatic v.2.8.4 (Bolger et al., 2014). HiSat2 v.2.1.0 (Kim et al., 2015) was used to map the reads to the *T. longispinosus* genome v.1 (GenBank accession: GCA_004794745.1_tlon_1.0; Kaur et al., 2019). We chose to use a single species genome as reference for all species to be able to later directly compare expression patterns between slave-making ants and their hosts as well as between queens and workers across species, with species as replicates. The counts table was created with HTSeq (Anders et al., 2015). To prevent spurious results due to low read counts, we removed from the counts matrix genes with less than 10 reads in at least four samples before the subsequent differential gene expression analysis with DESeq2 (Love et al., 2014). We started with the full model ∼Caste+Lifestyle+Caste:Lifestyle. The interaction turned out to be nonsignificant (no differentially expressed genes for the interaction), and we thus based all follow-up analyses on the two main factors only. All p-values were adjusted by false discovery rate (FDR) correction as implemented in DESeq2. To determine whether the significant slave-maker differentially expressed genes (DEGs) are more numerous than we would expect by chance, we ran 1000 permutations on the DEGs analysis in R.

To gain a deeper understanding for the functionality of DEGs, we 1) conducted a functional enrichment analysis, 2) inferred pathways in which the DEGs are involved, and 3) used a word mining approach based on longevity and fecundity terms. In detail, we ran Interproscan v.5.39-77.0 (Jones et al., 2014) locally to obtain GO information using the *T. longispinosus* predicted proteome (GenBank accession: GCA_004794745.1_tlon_1.0; Kaur et al., 2019) as query. The GO enrichment analysis was performed with the R package TopGO (Alexa & Rahnenführer, 2016), using the ‘parentchild’ algorithm and the Fishers exact test for significance. Furthermore, the proteome was annotated with KEGG functional ortholog numbers using the BlastKOALA web utility (last accessed: 11.02.2021; Kanehisa et al., 2016) with ‘Eukaryotes, Animals’ specified as Taxonomy group. KEGG pathway affiliation of genes that were differentially expressed in each of the four groups (queens, workers, slave-makers and hosts) was assessed using the online utility of KEGG mapper with *Apis mellifera* as reference species (Kanehisa & Sato, 2020), and visualized including the log2-fold change using pathview (Luo & Brouwer, 2013) in R v.3.6.3 (R Core team, 2020). KEGG pathway enrichment was assessed with the enrichKEGG function implemented in clusterProfiler v.3.14.3 (Yu et al., 2012) where the universe was defined as the total set of *T. longispinosus* protein predictions with a KEGG functional ortholog annotation and organism ‘ko’. For the text mining approach, we used gene annotations based on a BlastP search of the proteome versus the RefSeq invertebrate database, as query for a UniProt search. More specifically, we extracted the gene function information for each gene with entries from *C. elegans, D. melanogaster, A. mellifera* and searched for terms related to fecundity and longevity (Supplement Table S3), both traits which we assumed to differ between queens and workers (script available from Negroni et al., 2021). We conducted an enrichment analysis by conducting a Fisher’s exact test to test whether the number of genes with terms in fecundity or longevity within differentially expressed genes was higher than expected with respect to the complete proteome. The online tool Venny v.2.0.2 was used to generate the Venn diagram (https://bioinfogp.cnb.csic.es/tools/venny).

### Network analysis

To identify networks of co-expressed genes (modules), we constructed a weighted gene co-expression network analysis using the WGCNA (Langenfelder & Horwath, 2008) package in R. We used all genes that had passed the quality filtering step for the expression analysis (N = 8,327), thus not only the differentially expressed genes. Gene counts were normalized using the *varianceStabilizingTransformation* function from DESeq2 (Love et al., 2014). Following the WGCNA guidelines, we picked a soft-thresholding power of 8 for adjacency calculation. To associate modules to either caste or lifestyle, we first calculated the modules’ eigengene using the *moduleEigengenes* function and tested for module trait correlation using the *corPvalueStudent* function. The hub gene of each module (i.e., the gene with the highest connectivity within a module) was determined using the *chooseTopHubInEachModule* function. Moreover, for modules associated with caste or lifestyle, we tested whether the connectivity of genes with caste-specific expression and genes differed from the connectivity of those that were not differentially expressed. We ran linear models with connectivity as response and caste-specificity (queen, worker, no expression difference) as explanatory variable in R.

To determine the relevance of fecundity and longevity associated genes in lifestyle and caste associated modules, we downloaded a list of 123 genes that were assembled based on information from *Drosophila* and are part of the TI-J-LiFe pathways (TOR/IIS-JH-Lifespan and Fecundity) (Korb et al. 2021). We used the Flybase identifiers to download the corresponding *Drosophila* sequences and used a BlastX search to identify *T. longispinosus* proteins with a blast hit e-value < e-10. In cases where we had two or three hits, we selected the protein with the longest match, which mostly corresponded to the lowest e-value. In cases where we had >3 blast hits, we created a sequence alignment using MACSE v.2.03 (Ranwez et al., 2011). A Maximum Likelihood phylogenetic tree with 1000 bootstrap replicates was constructed with RAxML with PROTGAMMAWAG as protein substitution model. Based on the tree topology we chose the *T. longispinosus* sequence with the closest relationship to the target *Drosophila* sequence.

### Selected genes analysis

For this analysis, we added transcriptomes of four taxa that are not known to be parasitized by slave-making ants, which acted as a biological control for comparisons of selection intensity between slave-makers and hosts: as outgroup, two populations of *Cardiocondyla obscurior*, and the two non-host species of the *Formicoxenus*group, *Temnothorax nylanderi* and *T. rugatulus* (Supplement Table S1). We used Trinity v.2.8.6 (Grabherr et al., 2011) with standard settings to construct species-specific transcriptomes using both worker and queen transcripts. Nucleotide sequences were translated into amino acid sequences with TransDecoder (https://github.com/TransDecoder). As *de novo* transcriptomes are known to contain many transcript fragments as well as isoforms, we constructed orthogroups across all species with OrthoFinder (Emms & Kelly, 2015), including protein sequences derived from the *Temnothorax longispinosus* genome (Kaur et al., 2019), and retained orthogroups with a tlon-v1 ortholog only. After orthogroup construction, the *T. longispinosus* protein sequences derived from the genome were removed from the orthogroups from all downstream summary statistics and analyses. Since we only obtained few single copy orthologs, we used an inhouse script (Supplement S_script) to retain only a single sequence per species. In short, based on the pair-wise blast results from OrthoFinder the sequence with the highest sum of bit scores (i.e. best match to all other sequences) within each orthogroup was chosen as “centroid”. From each species the sequence with the best match to the centroid was chosen to create the single copy orthogroup. We used Clustal-omega v.1.2.4 (Goujon et al., 2010) to construct sequence alignments for each orthogroup, which were trimmed with TrimAL v.1.4.1 (Capella-Gutierrez et al., 2009) and the following settings: - gappyout -resoverlap 0.75 -seqoverlap 75 -backtrans. To test for signatures of positive selection we used the codeml implementation in ete3 v.3.1.1 (Huerta-Cepas et al., 2016) running the branch site test of selection as follows: ete3 evol --models bsA bsA1 --tests bsA, bsA1 --leaves --internals. The cluster-specific tree topology as inferred by RAxML v.8.2.12 (Stamatakis, 2014) was used as input tree, and each species was coded as foreground branch consecutively. Finally, we blasted the *T. longispinosus* sequence of each ortholog cluster with signatures of selection versus the *T. longispinosus* genome and only retained ortholog clusters with sequences mapping to a single location, as further means of preventing putative paralogs. We further ran a GO enrichment analysis to test for overrepresented functions among the selected genes (details above), and used the KEGG annotations to investigate the pathways in which genes with signature of selection are involved.

## Results

We set out to investigate the effect of lifestyle and caste on gene expression patterns across 15 different species of ants contrasting slave-making ants with their hosts, and queens with workers. We were mainly interested in determining whether there are common toolsets of genes characteristic for a specific lifestyle or caste. We additionally tested for genes under selection in the above-mentioned slave-maker-host pairs plus four additional non-host outgroup species.

In general, gene expression patterns seem to be most similar amongst phylogenetically close species. The principal component analysis revealed a very strong phylogenetic effect with 71% of variance explained by PC1, which separated the *Harpagoxenus / Leptothorax* group from the *Temnothorax* species (Supplement Figure S1), while PC2, explaining 8% of the variance, separated queens from workers. Caste had a much stronger effect on gene expression compared to lifestyle, as also evident in the heatmap dendrogram, where samples cluster according to caste rather than lifestyle (Figure 1). As our analysis did not reveal an interaction between lifestyle and caste (0 DEGs associated to the interaction term), we present the gene expression plus gene network results according to the main effects, caste and lifestyle, and finally the results from the selection analysis.

**Figure 1:**
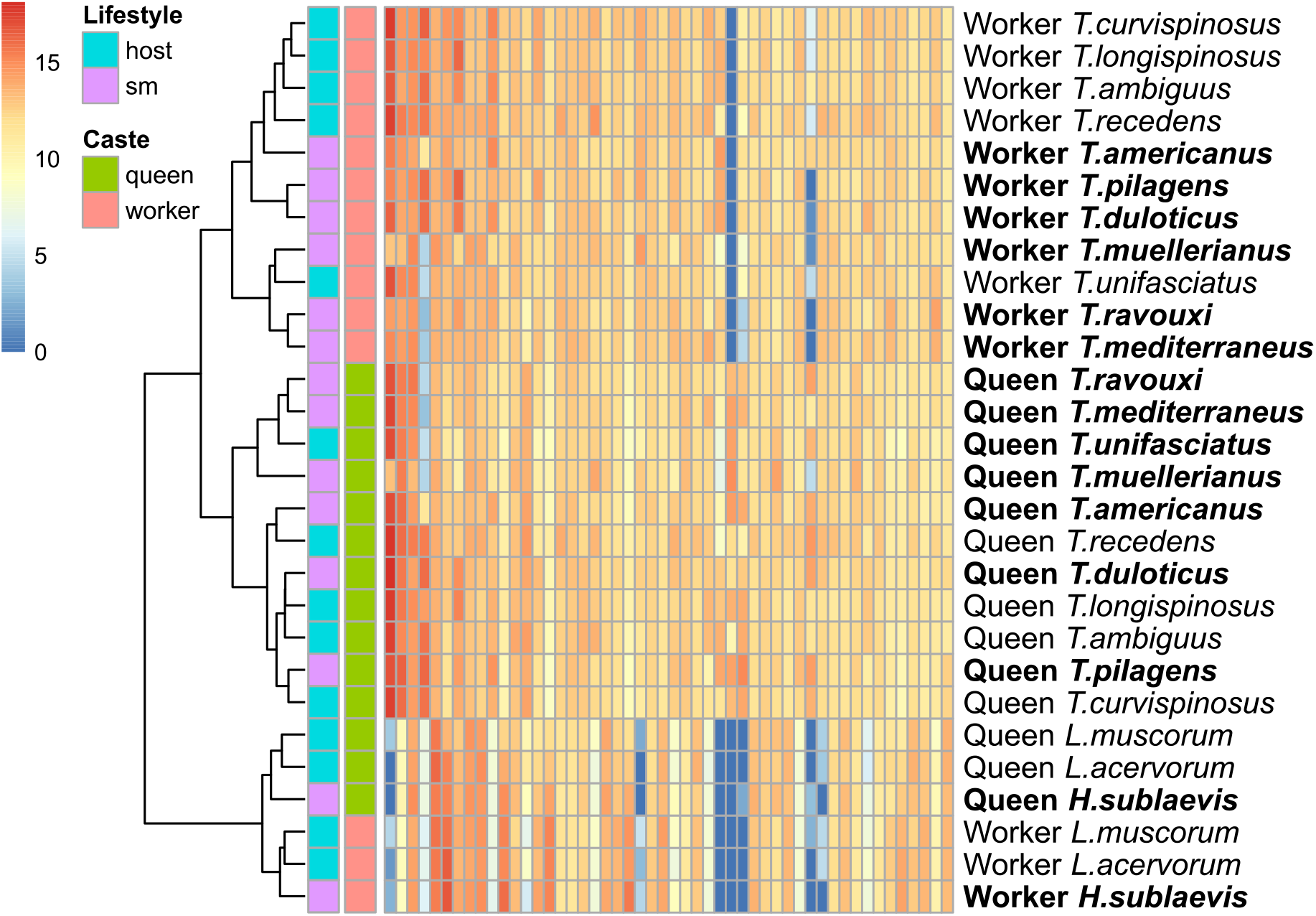
Sample dendrogram and heatmap depicting a strong clustering according to caste and a weaker effect of lifestyle. The heatmap is based on the top 50 highest expressed genes. Slave-making species are highlighted in bold.

### Lifestyle

We detected 62 differentially expressed genes (DEGs) between lifestyles (host versus slave-maker species) cumulative across the five origins of slavery (see Supplement Table 1 DEGs). A permutation test with 1000 iterations revealed that this is more than expected by chance (p = 0.04). Six genes were consistently higher expressed in all seven slave-maker species compared to hosts, of which three were annotated: “protein DVR-1 homolog”, “sulfotransferase family cytosolic 1B member 1-like”, and “ATP-dependent DNA helicase II subunit 1-like”. Only a single gene, “NADPH oxidase 5”, showed higher expression in all seven host species compared to slave-makers (Figure 2). Various metabolic processes were significantly enriched among the differentially expressed genes between lifestyles (total = 21 processes). Among genes up-regulated in slave-makers, 14 processes were enriched, and three functions were enriched in genes up-regulated in hosts, including “cell death” (Supplement_GO). In addition to the functional level, we also investigated whether the genes in the lifestyle DEGs set were overrepresented in specific pathways. Due to the low number of lifestyle-associated DEGs, and only about half of these with KEGG annotation, the enrichment analyses for the complete set of lifestyle DEGs, and the separate host and slave-maker gene-sets, did not result in any overrepresented functions.

**Figure 2:**
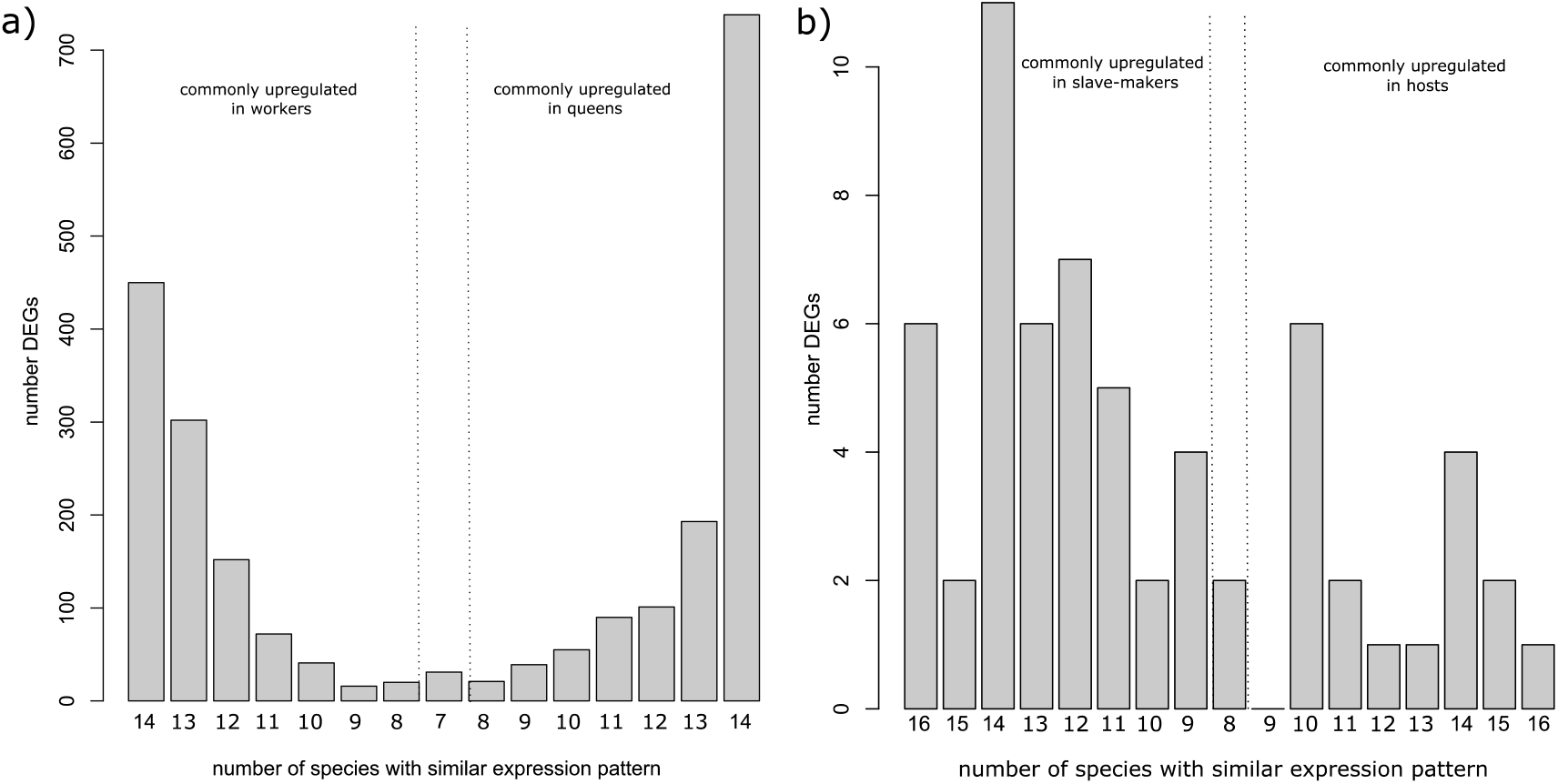
Number of differentially expressed genes (DEGs) which showed shared expression patterns across a majority of study species during the pairwise comparisons of queens vs workers or slave-makers vs hosts. a) Number of DEGs upregulated in workers (left) or upregulated in queens (right) across more than half the study species. b) Number of DEGs upregulated in slave-makers (left) or upregulated in hosts (right) across more than half the study species. The vertical lines delimit the start of a majority of species in either direction, with the bar between the lines showing DEGs that showed a common expression pattern in one direction in half the species, and in the other direction in half the species -e.g. in graph a), DEGs which were upregulated in queens in 7 species, and upregulated in workers in the other 7 species. Please note that the host-slave-maker pairs add up to 16 comparisons instead of 14, since *Harpagoxenus sublaevis* parasitizes two host species: *L. acervorum* and *L. muscorum*.

Host colonies are generally larger than slave-maker colonies, *i*.*e*., host queens are more fecund than slave-maker queens. Moreover, fecundity is often positively correlated with longevity in social insects (Korb & Heinze, 2016; Negroni et al., 2016). We therefore conducted a word mining approach based on UniProt entries to investigate whether genes with known fecundity or longevity functionality were overrepresented in the sets of DEGs between hosts and slave-makers (Table 1). We indeed recovered more DEGs putatively associated with fecundity than expected by chance (N = 31; Fisher’s exact test, p-value = 6.619e-09), but not for the longevity associated genes (Table 1).

**Table 1:** Number of differentially expressed genes with a fecundity or longevity functionality based on a text mining approach using the UniProt database as a reference. Results of Fisher’s exact test are given indicating whether the number of genes with either fecundity or longevity association in caste and lifestyle are more than expected by chance.

| *Treatment* | **Fecundity** |  | **Longevity** |  |
| --- | --- | --- | --- | --- |
|  | N of genes | P-Value | N of genes | P-value |
| ***Caste*** | **1076** | **2.2e-16** | **412** | **0.0003** |
| *Queen* | 630 |  | 262 |  |
| *Worker* | 447 |  | 150 |  |
| ***Lifestyle*** | **31** | **6.619e-09** | **13** | **0.39** |
| *Host* | 9 |  | 6 |  |
| *Slave-maker* | 23 |  | 7 |  |

We obtained 10 modules of co-expressed genes (named after colours as given by the *WGCNA* package), of which one module, red, was positively associated with the slave-making lifestyle (Figure 5). This module contained 191 genes, of which 19 were up-regulated in slave-makers and none in hosts. Genes associated with this lifestyle module were enriched for “reactive oxygen species metabolic processes”, “oxidation reduction processes”, or “fatty acid biosynthetic process” (Supplement_WGCNA). Genes up-regulated in slave-makers had the highest connectivity in this module associated with lifestyle (Supplement Figure S2).

### Caste

Caste had a much stronger effect on gene expression than lifestyle (χ^2^= 2496.8, df = 1, p-value < 2.2e-16), with 2,321 differentially expressed genes (DEGs) between queens and workers. Around half of all caste-specific DEGs showed consistently higher expression in queens (N=738 out of 1,295) or in workers (N = 450 out of 1,026) across all species. For example, genes for an *insulin-like growth factor 2, mRNA-binding protein 1, maternal protein exuperantia, histone deacetylases*, and several *serine/threonine-protein kinases* were more highly expressed in queens of all 15 species, while *pro-corazonine like* had higher counts in workers of all but one species. The number of genes with consistent expression across taxa was significantly lower for the lifestyle comparison (χ^2^= 36.865, df = 1, p-value = 1.266e-09).

As queens and workers strongly differ in fecundity and longevity, we used a word mining approach based on UniProt entries to investigate whether genes with known fecundity or longevity functionality were overrepresented in the sets of caste-specific DEGs (Table 1). Indeed, we detected more fecundity-associated terms (N = 1076; Fisher’s exact test, p-value = 2.2e-16), and more longevity-associated terms (N = 412; Fisher’s exact test, p-value < 0.0003) than expected by chance.

In the caste-specific DEG set, 39 gene ontology (GO) functions were significantly overrepresented, many of which were associated with metabolic and biosynthetic processes (Supplement_GO). Genes up-regulated in queens were enriched for 60 functions linked to various metabolic processes or stress responses. Among the worker up-regulated genes, 78 functions were significantly enriched, also belonging to metabolic and biosynthetic processes, but also oxidation-reduction. In addition to the functional enrichment, we conducted a KEGG-pathway enrichment analysis. In total, we found 27 pathways enriched amongst the caste-specific DEGs, 12 for genes up-regulated in queens, and 61 genes up-regulated in workers (Supplement_KEGG). Multiple putatively reproduction-associated and repair pathways, such as “meiosis-yeast”, “cell cycle”, “DNA replication”, “RNA transport,” or “ribosome biogenesis,” were enriched in queens, and pathways such as “olfactory transduction” and “longevity regulating pathway” (Figure 3) were enriched in genes up-regulated in workers (Supplement_KEGG).

**Figure 3:**
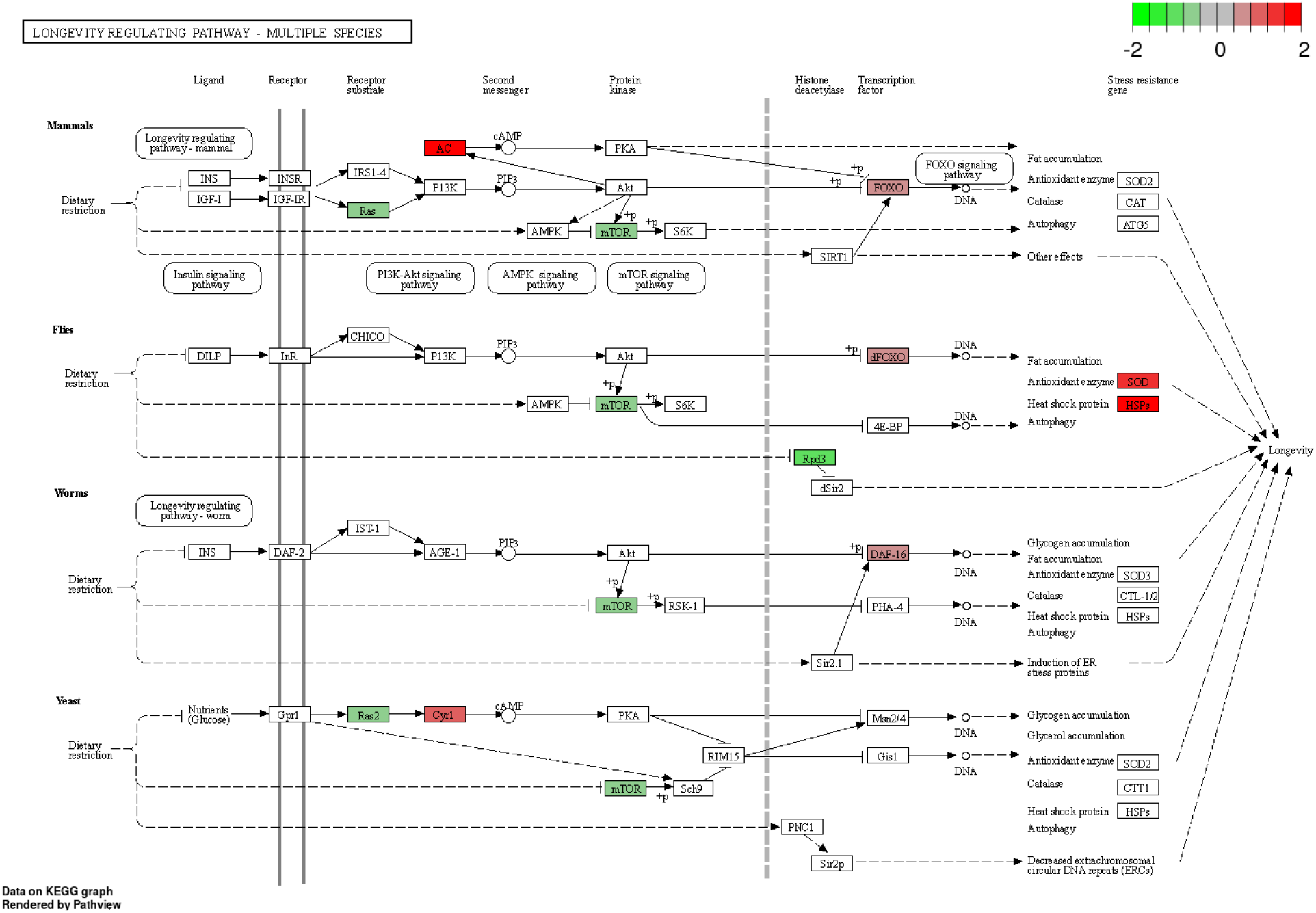
KEGG map depicting the “longevity regulating pathway” with up-regulated genes in queens (negative logFC; green) versus up-regulated in workers (positive logFC; red).

The gene network analysis resulted in five modules associated with caste, three of which were positively associated with queens (yellow, green, magenta), and two with workers (black and blue) (Figure 5). There was a trend for the pink module to also be associated with caste (p = 0.07), and it was included in the following analyses (for details see: Supplement_WGCNA). In all three worker associated modules (black, blue, pink), connectivity was highest for worker up-regulated genes, lowest for queen up-regulated genes, and intermediate for genes that were not differentially expressed. The same pattern was observed in queen-specific modules (yellow, green, magenta), in which the queen up-regulated genes had the highest connectivity (Supplement_WGCNA).

### Comparison to other species

We identified 84 genes associated with one of the 10 WGCNA modules that have also previously been linked to fecundity or longevity in Drosophila. We obtained these from the “TI-J-LiFe” list containing 123 candidate genes (Korb et al., 2021). About half of these (N = 60) were found in the turquoise module, which was neither associated to caste nor lifestyle, and eleven were found in the blue, caste-linked module (Supplement_WGCNA). To determine whether overrepresented gene functions linked to caste associated modules from our Formicoxenus-group data set could be extrapolated to other ants or even termites, we compared our modules to Morandin et al. (2016) and Lin et al. (2021). Only three functions were shared between enriched GO-terms of caste associated modules in our slave-making data set (N = 46) and enriched GO-functions in Formica caste associated modules (N = 155; Morandin et al., 2016), namely “cellular protein modification process”, “protein modification process”, “monovalent inorganic cation transport” (Table 2). Eight terms were shared with caste associated modules in termites (N = 277; Lin et al., 2021) (Table 2). No terms were shared amongst the three studies. However, these results should be taken with caution as most enrichment algorithms take the hierarchical structure of GO terms into account. Thus one may have a lower or higher term in the hierarchy, which does not mean they are essentially different.

**Table 2:**
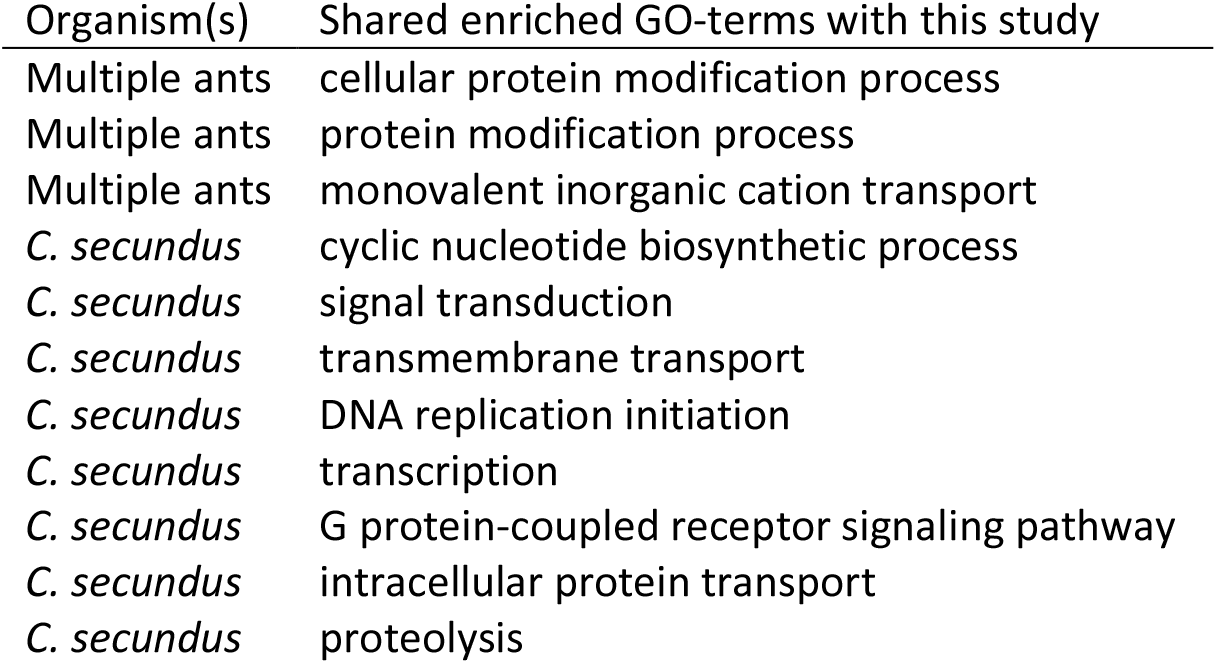
Shared enriched GO terms between queen-worker caste associated modules from this study, a study on 16 ant species from multiple genera (Morandin et al. 2016), and the termite *Cryptotermes secundus* (Lin et al. 2021).

As queen and workers differ in fecundity and longevity, we investigated whether the same set of fecundity-longevity associated genes of the “TI-J-LiFe” list (N = 84) which were found in the termites (Lin et al., 2021) could be found in our data set. There were two genes, Ras64B and Kr-h1, associated with a queen-worker co-expression module in termites and in Crematogastrini.

### Selected genes analysis

The transcriptomes of the 19 taxa, including four samples of non-host species, consisted of 67,150-155,629 transcripts with 32,584-51,743 open reading frames and 92-96% DOGMA completeness. In total, we obtained 10,699 ortholog clusters of which 5,826 clusters contained at least one transcript per species. 1,398 clusters remained after trimming and filtering for single copies per species. We identified 660 signatures of selection across all species, that are associated with 424 ortholog clusters, and mapped to a single location on the genome (Supplement_codeml). Between 6-92 genes showed signatures of selection in each species (Supplement_codeml), but there was no difference in the number of selected genes between lifestyles (quasipoisson model, X^2^ = 1.33, d.f. = 2, p = 0.51). In slave-makers, genes with signatures of selection were significantly enriched in functions such as protein modification, demethylation and energy maintenance (Figure 4a, Supplement_codeml). In hosts and non-hosts, several regulatory and metabolic functions are significantly overrepresented in genes with signatures of selection (Figure 4b+c).

**Figure 4:**
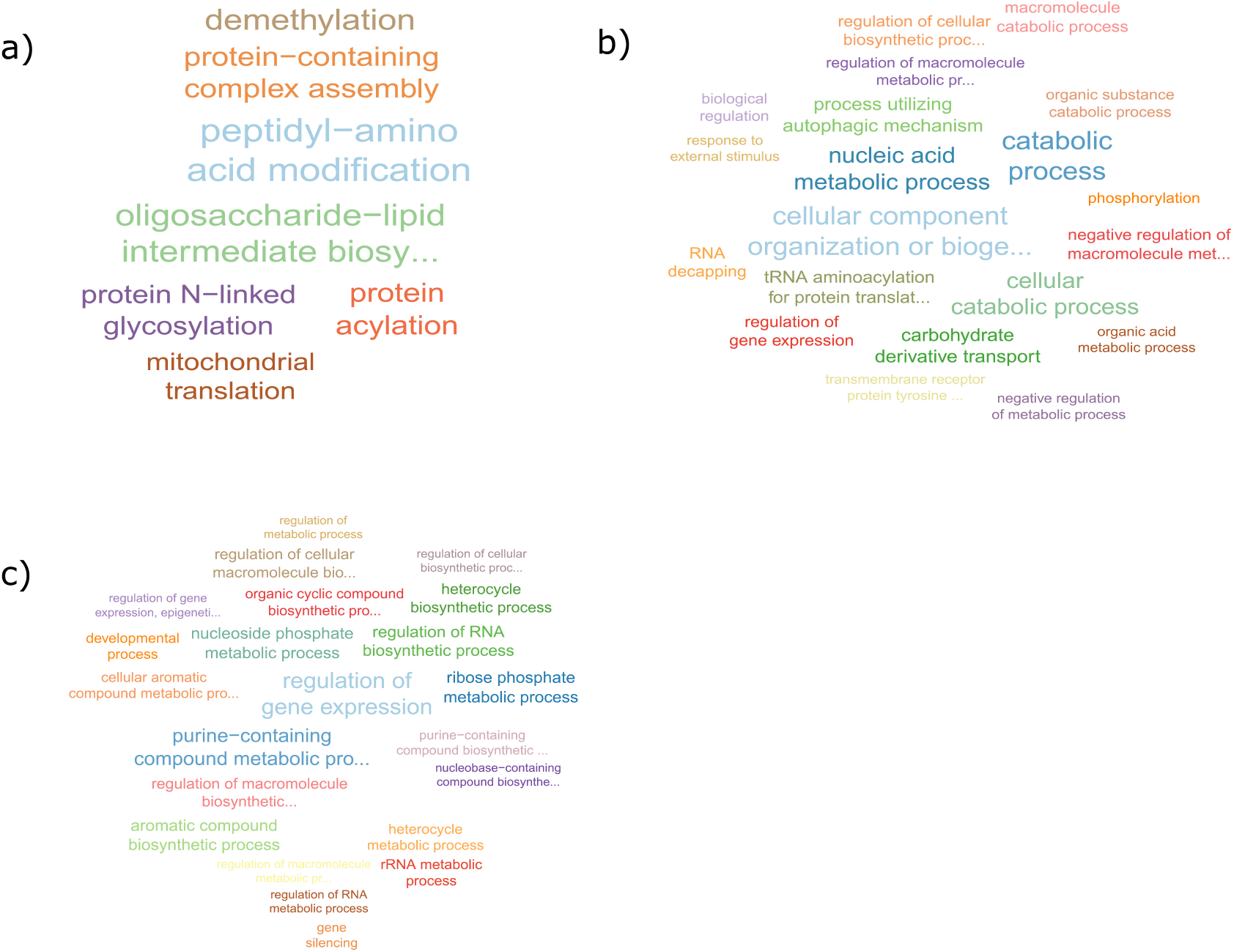
Word clouds of enriched GO functions based on genes with signature of selection in a) slave-makers, b) hosts, c) non-hosts.

**Figure 5:**
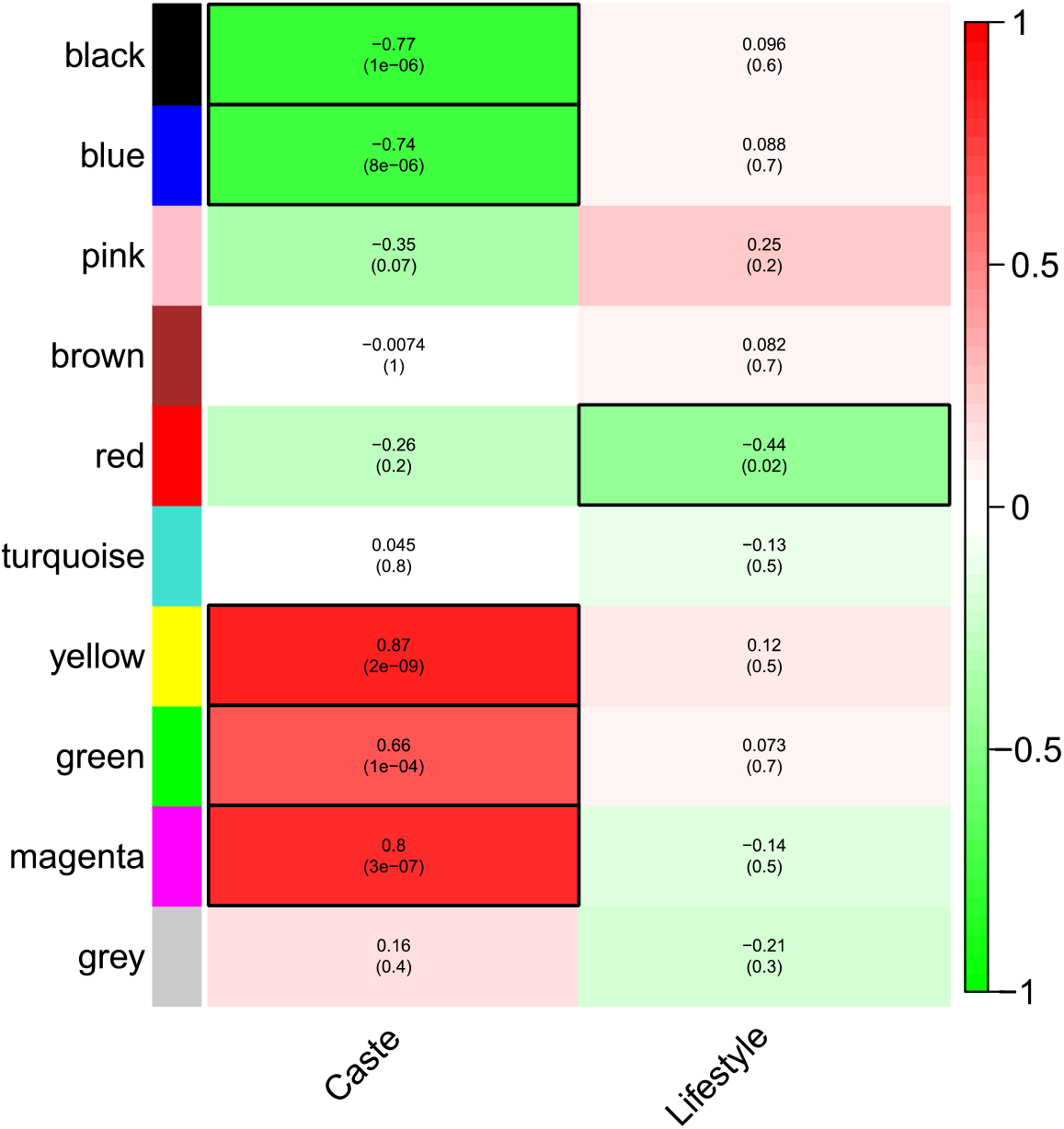
Module trait relationship of the 10 gene co-expression clusters with caste and lifestyle. The strength of the correlation is given in the upper numbers and significance levels in parentheses below. Significant modules are highlighted by black boxes. Modules are labelled by colours according to standard WGCNA output. Green indicates modules of genes overexpressed in workers / slave-makers, red indicates those with genes more expressed in queens / hosts.

We identified 114 genes that were both differentially expressed and additionally showed signatures of selection (Supplement_overlap). Of these, a single gene was differentially expressed between castes and lifestyles and positively selected in *T. unifasciatus* (TLON_06615-RA; pleckstrin homology domain-containing family F member 2 isoform X1). All remaining 113 were differentially expressed only between castes. 35 of these caste-specific DEGs showed signatures of selection in hosts, 23 in non-hosts, and 57 in slave-makers. Interestingly, many nucleotide metabolism pathways were included in the overlapping gene lists of differentially expressed and selected genes, including purine, alanine, aspartate, valine, and the “mTOR signalling pathway” (Supplement_overlap).

## Discussion

Eusocial insects are characterised by their sophisticated division of labour, which led to the evolution of different castes. Ant queens are the main reproductives, known for their long life of up to several decades and high fecundity (Keller & Genoud, 1997), while the mostly infertile and short-lived workers take care of the brood, foraging, and nest defence (Hölldobler & Wilson, 1990). In slave-making species, however, workers do not perform the “general” worker chores. They raid host nests in summer and often are permitted to lay male-destined eggs in the presence of the queen. Our study shows that whole-body transcriptomes of seven species pairs of the “*Formicoxenus* group” of Crematogastrini (Blaimer et al., 2018; *Harpagoxenus, Leptothorax*, and *Temnothorax*) differ considerably more between castes (queen vs. worker) than with lifestyles (slave-maker vs. non-parasitic species). Caste polyphenism is based on differential gene expression, and our transcriptome analyses show that the expression of 1,188 genes consistently differs between queens and workers of all studied species, regardless of lifestyle. Although the evolution of a parasitic lifestyle is associated with considerable changes in morphology and behaviour, only few transcriptomic shifts from host to slave-making species were consistent across the five independent origins of slave-making.

Approximately 40x more genes were differentially expressed between castes than between lifestyles (2,321 vs. 62), and six out of ten co-expression modules were caste-associated in contrast to a single lifestyle module. This difference might in part reflect the single evolutionary origin of caste diphenism in ants, whereas slavery evolved repeatedly (Beibl et al., 2005; Feldmeyer et al., 2017; Prebus, 2017). Nevertheless, as slave-making workers are often fertile and do not take over normal worker chores, we had expected to find a considerable interaction between caste and lifestyle in gene expression as well as in gene connectivity, as caste differences are less pronounced in slave-making species. The lack of the interaction indicates that slave-maker and host queens and workers are rather similar on a molecular level. Furthermore, it corroborates the result of a previous study where caste differences also exceeded differences due to other traits, such as worker sterility, queen number, or invasiveness (Morandin et al., 2016).

While in this study, half of the genes that were differentially expressed between queens and workers show the same expression pattern across all species, another study identified only a single gene (the myosin light chain) similarly expressed between queens and workers a set of 16 species of multiple genera (Morandin et al., 2016). The reason for this discrepancy may be explained by the different species relationships as well as the underlying data basis. We studied species within a single, closely related clade and used the genome of a single species as reference. In contrast, Morandin et al. (2016) investigated species from five genera in different subfamilies based on de novo assembled transcriptomes. Additionally, there is evidence for similarities in pathways across different lineages of social insects (ants, bees, wasps) rather than a “common toolkit” of genes responsible for the caste phenotype (Berens et al., 2015). In our phylogenetically more restricted data set however, we identified 1,188 genes representing the “core set” of queen-worker differences across the 15 species. For example, the gene “*corazonin*” has been shown to control social behaviour and caste identity in ants (Gospocic et al., 2017). The gene “*maternal protein exuperantia*”, a maternal effect gene which is needed for proper localisation of the *bicoid* RNA during oocye formation (de Olivereira et al., 2017; McDonald et al., 1991), but also plays a role in *Drosophila* spermatogenesis (Hazelrigg et al., 1990) was up-regulated in all queens. Also, “G1/S-specific cyclin-D2” (cycD) was more strongly expressed in queens compared to workers in all but one species. This gene is involved in cell cycle regulation and part of the “FoxO signalling” and “Wnt signalling pathway”, both important regulators of longevity. It could thus be associated with the lifespan differences between these two castes. More generally, half of the genes differentially expressed between caste were associated with the UniProt functionalities “fecundity” and “longevity.”

Reflecting the convergent evolution of ant slavery (Beibl et al., 2005; Feldmeyer et al., 2017; Prebus, 2017), the different parasitic species not only show pronounced differences in their morphology, but also in raiding behaviour (Brandt et al., 2006; Johnson, 2008; Kleeberg & Foitzik, 2016). Species-specific raiding patterns are mirrored by species-specific gene expression patterns (Alleman et al., 2018) and genes under selection (Feldmeyer et al., 2017). Gene expression differences between pairs of slave-makers and their hosts showed much more variation across species (representing different origins of slaver-making) than the differences between queens and workers, *i*.*e*., there are many more idiosyncrasies in how slave-makers and hosts differ. Nevertheless, we found 62 genes that varied with lifestyle, including genes for fatty acid synthases. This could be indicative either of differences in fat synthesis and maybe storage between the two lifestyles, but these synthases may also be involved in the synthesis of cuticular hydrocarbons which are used as communication signals to discriminate species and castes (Leonhard et al., 2016).

As slave-makers are the derived lifestyle with species-specific behaviours and traits, we expected to find more genes under selection in slave-makers than in hosts and/or non-host species, however the number of selected genes did not differ between lifestyles. Genes with signatures of selection were significantly enriched in functions such as protein modification, demethylation, and energy maintenance. An interesting candidate among the genes that showed signatures of selection in only one or a few of the species is venom protease-like in *T. muellerianus*. Though at present nothing is known on the composition of the venom used by *T. muellerianus* to kill host ants during slave-raids, many Hymenopteran venoms are proteases (Touchard et al., 2016). On a higher level, many metabolism pathways were included in this overlapping gene list, including purine, alanine, aspartate, valine, to name just a few, and the “mTOR signalling pathway”.

## Conclusion

Social parasitism represents a derived state with clear differences to host and non-host species from morphology to behaviour. In most social parasites, queens and workers are more similar than in host species. Examining gene expression patterns and gene regulatory networks of 15 different ant species spanning five origins of slave-making, we expected to find an interaction between caste and lifestyle effects on gene expression patterns. Despite the phenotypic differences, between slavemaker and host castes, gene expression profiles were remarkably similar. Our study corroborates previous results indirectly indicating species-specific expression and selection patterns with respect to lifestyle, thus little common difference between host and slave-maker species. However, we observed a very strong and reliable effect of caste on gene expression. Within our broad taxonomic species spectrum, we were able to identify a core-set of 1,188 caste-specific genes, which show a consistent expression pattern across all species irrespective of lifestyle, pointing to a “genetic toolkit” in this set of related ant species.

## Acknowledgements

We would like to thank Tilman Shell for writing the ortholog cluster filter script for us. This work was supported by the Deutsche Forschungsgemeinschaft (grant numbers Bo 2544/12-1 to EBB, Fo 298/20 to FO, He 1623/40 to JH).

## Data Accessibility

All raw sequence data underlying this study have been deposited in the National Centre for Biotechnological Information (NCBI) Sequence Read Archive (SRA) and will be accessible upon publication of this manuscript (BioProject accession number XY will follow in the next version of the manuscript).

## Author Contributions

The study was conceived by JH, EBB and SF, and was designed by EJ, BF, JH, EBB and SF. DG provided ant samples. BF, CS, JW and EJ conducted the analyses. All authors contributed to writing the paper. Authors declare no conflict of interest.

